# CICADA: A unified framework for NWB-based neurophysiological data analysis

**DOI:** 10.64898/2026.07.03.736318

**Authors:** Myriam Hamon, Jules Lebert, Julien Denis, Caroline Filippi, Anthony Renard, Pol Bech, Mauro Pulin, Axel Bisi, Daniel Molinuevo Gomez, James B. Priestley, Sylvain Crochet, Carl C. H. Petersen, Rosa Cossart, Michel A. Picardo, Robin F. Dard

## Abstract

Neurophysiology datasets are becoming increasingly complex, combining behavioral measurements with high-dimensional neuronal activity recordings coming from optical and/or electrophysiological measurements. The Neurodata Without Borders (NWB) standard has emerged in the community as the format of record. While standardized and widely used preprocessing tools generating NWB files have been developed, extensible frameworks for scientific analysis downstream of the NWB ecosystem are still under-represented. We present CICADA, a Python framework dedicated to analysis of neurophysiological data in the standardized NWB format. The toolbox is built as three hierarchically-organized packages: ***cicada-nwb*** (NWB access layer), ***cicada-analysis*** (plugin-based analysis engine and tool library), and ***cicada-gui*** (PyQt5 desktop application at the head of the pipeline). **Beyond this architectural separation, CICADA is built around a central design principle: supporting a continuum from turnkey use to community extension**. Researchers can use the complete GUI-driven ***cicada-gui*** workflow without writing code, programmatically use existing analysis plugins from ***cicada-analysis***, or extend the framework with their own analysis plugins, written with the ***cicada-nwb*** access layer and the *cicada-tools* library and shared as lab packages that CICADA discovers automatically once installed. The same analysis plugin runs identically in interactive GUI and parameter-configured headless modes, enabling reproducible multi-session, multi-animal group analyses. We illustrate the versatility of CICADA with example analyses of behavioral, calcium imaging and extracellular electrophysiology. CICADA is open source, actively maintained, and designed so that any laboratory can share its analyses as a lab package, without modifying the core framework.

## INTRODUCTION

Advances in neurophysiological recordings, including two-photon (2p) and widefield calcium imaging, fiber photometry and extracellular electrophysiology, now enable recordings of larger and larger populations of neurons in behaving animals, thus generating terabyte-scale datasets per laboratory each year. In order to publish and share these datasets, The Neurodata Without Borders (NWB) format has become a standard in the community, with currently more than 1,200 public datasets and more than 2,200 users as of November 2026 on DANDI archive (“DANDI,” n.d.; Rübel et al., 2022).

Simultaneously, the number of NWB compatible tools has grown over the past years. These existing tools mainly cover data preprocessing, such as CaImAn (Giovannucci et al., 2019), OptiNiSt (Yamane et al., 2025), and Suite2p (Pachitariu et al., 2017) for cell detection, fluorescence trace extraction and spike estimates in imaging data; while SpyKING circus (Yger et al., 2018) can be used as a spike sorting algorithm starting from NWB files, and other spikesorters (for example Kilosort 1-4 (Pachitariu et al., 2024) or MountainSort (Chung et al., 2017)) as well as other processing steps can be done starting from NWB files through SpikeInterface (Buccino et al., 2020), a framework for extracellular electrophysiological recordings analysis. Some tools have been developed to facilitate NWB conversion such as NeuroConv (Mayorquin et al., 2025) and NWB GUIDE (Flynn et al., 2026); others allow for NWB exploration such as NWB explorer (“Open Source Brain,” n.d.) or NWB Widgets (“NWBWidgets,” n.d.). The development of a large number of NWB compatible tools for neurophysiology data preprocessing accelerated the adoption of the format by many laboratories but a standard framework for downstream scientific analyses was often lacking following data preprocessing and data conversion.

In practice, lab-specific analysis scripts are typically under-documented, parameter-laden, and hard to share, making it an obstacle for reproducible research. Facing this issue, the community is progressively developing such analytical tools. We released the GUI-based NWB-native analysis pipeline CICADA (Calcium Imaging Completely Automated Data Analysis) (“Cossart Lab / CICADA,” 2019). This pipeline, primarily designed to analyze calcium imaging data, was used for various published analyses (Bech et al., 2026; Bocchio et al., 2020; Dard et al., 2022; Leprince et al., 2026, 2023; Ratsifandrihamanana et al., 2023). Since then, the list of NWB-based analysis tools has grown and other pipelines relying on the NWB format have been developed, such as Spyglass (Lee et al., 2024) and Pynapple (Viejo et al., 2023).

Here, we present an updated version of CICADA, a modular NWB-native analysis framework spanning interactive and headless workflows and open to community extension, designed to cover a wide range of user needs and coding skills. On one side of the spectrum, the pipeline offers the possibility to run ready-to-use analysis plugins, ranging from data visualization to complex multi-step workflows, across calcium imaging, widefield imaging, electrophysiology, and behavioral data, entirely through a Graphical User interface (GUI), or through a headless runner with minimal scripting. On the other, it lets developers turn their own analyses into plugins, built on the provided NWB access layer and analysis tools and shared through lab packages. To achieve this level of flexibility, the pipeline is organized into three modular packages: ***cicada-gui*** supporting the graphical interface, ***cicada-analysis*** the analysis engine and ***cicada-nwb*** a NWB reader package.

We present CICADA by starting with its intuitive and ready-to-use graphical interface, which requires no programming experience, and progressively introduce its core engine, from the plugins supporting data analysis to its innermost components that allow one to write new plugins. We first show the general architecture of the pipeline (results section 1, R1). We then describe ***cicada-gui***, the head of the pipeline, and ***cicada-analysis***, the underlying analysis engine (R2). We further provide examples of NWB-based data analysis, showing generalization of the pipeline across datasets not explicitly targeted by the pipeline design (R3). Finally, we show how laboratories extend the plugin collection with lab packages, building on ***cicada-nwb*** and *cicada-tools* and discovered through standard Python entry points (R4).

This ordering reflects the design principle: each layer is independently useful, adopting the framework is not an all-or-nothing commitment and users can decide where to onboard from a GUI-driven workflow to writing and sharing their own plugins. CICADA provides a platform for researchers to analyze their NWB data in a standardized and reproducible way, regardless of their coding skills.

## RESULTS

### R1. The three-package architecture

The NWB format provides a standardized, self-describing container for neurophysiology and behavioral data, and its adoption has accelerated the development of shared preprocessing tools and public datasets. Building directly on NWB rather than on lab-specific file formats, any NWB-compliant dataset can be analyzed without conversion or custom loading code, making it the natural choice as CICADA’s primary input format.

CICADA is organized as three-layered packages, ***cicada-gui, cicada-analysis***, and ***cicada-nwb***, arranged from a fully graphical no-code interface down to a bare NWB access layer (Figure 1A). Each package depends on the one below, but can be used as an entry point in its own right, without requiring the layers above. This design reflects a core principle: researchers can onboard at any level of the stack without committing to the full framework.

**Figure 1.**
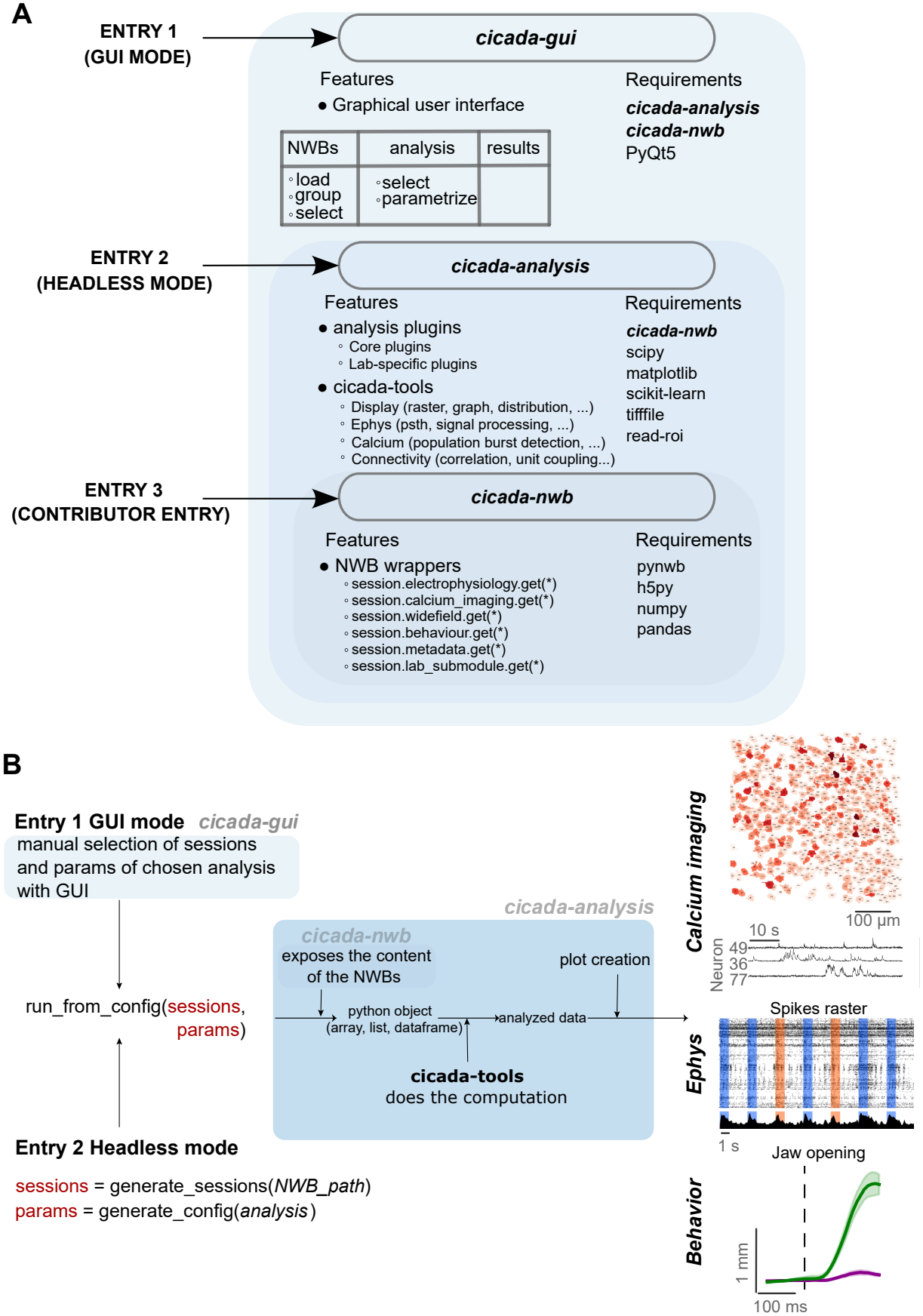
Architecture overview. **(A)** CICADA is composed of three packages, each representing an independent entry point into the framework. Entry 1 (GUI mode): *cicada-gui* is a PyQt5 desktop application that requires *cicada-analysis* and *cicada-nwb*, and provides a graphical interface for loading NWB sessions, selecting analyses, and parametrizing runs without writing code. Entry 2 (Headless mode): *cicada-analysis* provides the plugin-based analysis engine and the *cicada-tools* computation library, and requires only *cicada-nwb*; it can be used programmatically via *run_from_config()* with YAML-defined sessions and parameters. Entry 3 (Contributor entry): *cicada-nwb* is the base NWB access layer, exposing session data through thematic submodules (*session*.*electrophysiology, session*.*calcium_imaging, session*.*widefield, session*.*behavior, session*.*metadata, session*.*optogenetic, session*.*<lab>*) with a small set of scientific Python dependencies. **(B)** Execution workflow across entry points. In GUI mode (Entry 1), sessions and analysis parameters are selected interactively through *cicada-gui*, which calls *run_from_config*(sessions, params). In headless mode (Entry 2), sessions and parameters are generated programmatically via *generate_sessions*() and *generate_config*(), then passed to the same *run_from_config*(sessions, params) call, ensuring strict equivalence between the two modes. In both cases, *cicada-nwb* exposes NWB file contents as Python objects (arrays, lists, dataframes), *cicada-tools* performs the computation on these objects, and results are returned as analyzed data and figures. Example outputs are shown for three data modalities: calcium imaging (cell map and fluorescence traces), extracellular electrophysiology (spike rasters), and behavior (jaw movement trajectory).

At the head of the pipeline, ***cicada-gui*** is a PyQt5 desktop application that exposes the full analysis engine through a graphical interface requiring no coding. On launch, the GUI scans either a user-specified local folder for NWB files or a user-selected DANDI dataset with the choice to stream the data or download them. The GUI then presents the selected NWB files in a session browser in which each file maps to a unique session identifier, establishing a one-to-one correspondence between files and experimental sessions. Sessions can then be assembled into groups and saved or loaded as YAML files. An analysis tree browser, populated dynamically after NWB loading, organizes all discovered plugins from ***cicada-analysis*** into collapsible families. Selecting an analysis in the GUI tree renders its parameter form automatically, allowing for manual selection of all its parameters. Clicking RUN launches the ***cicada-analysis*** plugins using the selected sessions and parameters.

Underlying the GUI, ***cicada-analysis*** provides the plugin-based analysis engine. Its central abstraction is the *CicadaAnalysis* base class, which every analysis plugin inherits from and which defines three methods: *check_data(), setup_parameters()* and *run_analysis()*. New plugins are discovered through standard Python entry points by ***cicada-analysis*** without modifying any core framework code, and a single parameter schema drives analysis parametrization rendering in the GUI, YAML configuration, and headless execution, keeping interactive and non-interactive runs strictly equivalent. Bundled within ***cicada-analysis*** is *cicada-tools*, a standalone library of reusable analysis functions, more than 250 documented functions organized into 12 submodules. All ***cicada-analysis*** plugins delegate their core computation to *cicada-tools*, meaning these functions can also be imported directly in notebooks or custom scripts independently of the plugin system and of the NWB format.

At the base of the architecture, ***cicada-nwb*** is a lightweight NWB access layer with no dependency on the analysis engine or GUI. It has a single entry point, *NWBSession*(*path*), and provides a set of data access methods organized into thematic submodules: *session*.*calcium_imaging, session*.*electrophysiology, session*.*behavior, session*.*widefield, session*.*metadata, session*.*optogenetic* and lab submodules installed with lab packages (R4). Data retrieval uses lazy loading, meaning that only requested slices are loaded into memory, making it practical for the gigabyte-scale files routinely produced by modern neurophysiology experiments.

Together, these three packages define a continuum from turnkey use to community extension, in which each level is independently useful and adoption of a higher level of integration is always optional (Figure 1). A researcher may start with the complete ***cicada-gui*** workflow without writing code, move to the headless runner to script multi-session analyses and, when an analysis is missing, write a plugin on top of ***cicada-nwb*** and *cicada-tools* and share it as a lab package. The same analysis unit, the plugin, runs identically across all execution contexts.

### R2 - A. cicada-gui — session selection, analysis parametrization, execution

***cicada-gui*** is the recommended entry point for researchers who want to run existing analysis plugins without writing code. The GUI scans a user-specified folder for NWB files and allows the user to assemble them into named groups, saved as YAML files. Grouping is facilitated by a built-in *groupby*() utility that filters sessions based on NWB metadata fields, such as subject, session type, or experimenter, eliminating the need for manual file organization. These session YAMLs serve as the common currency across all entry points: a group defined in the GUI can be reused for session loading in the GUI or directly in headless mode without modification.

Once sessions are loaded, the analysis tree lists all plugins discovered by the engine, organized into families and filtered by data compatibility through each plugin’s *check_data*() method, ensuring that only analyses whose required data types are present in the loaded sessions are selectable. Selecting a plugin opens an interactive parametrization window, automatically rendered from the plugin’s *setup_parameters*() declaration, with no GUI-specific code required in the plugin itself. Parameters can be adjusted interactively and exported as a standalone YAML file, which can then be loaded again through the GUI or passed directly to *run_from_config*(sessions, params) for headless replay.

Clicking RUN launches execution in a worker thread, keeping the main interface responsive during execution, allowing users to launch multiple analyses simultaneously. Upon completion, a combined YAML encoding the session list, all parameter values, and package versions is written alongside results. This combined configuration file supports two reproduction paths: *run_from_config*(sessions, params), which allows sessions and parameters to be modified or reused independently, and *run_from_config*(combined), which replays a previous run exactly as a single self-contained file.

Full installation instructions, a quickstart guide, and annotated screenshots are available in the online documentation (https://cicada-analysis.readthedocs.io/en/latest/gui/).

### R2 – B. cicada-analysis — no-GUI execution of analysis plugins

For users who want to run analyses without using the GUI, ***cicada-analysis*** supports fully headless execution via *run_from_config*(), with strict equivalence to the GUI workflow (Figure 2). This enables analyses to be deployed on remote servers or computing clusters, removing the dependency on a local machine and display server. Starting from a folder of NWB files, *generate_sessions*() produces a session YAML and *generate_config*() produces a parameter YAML file pre-populated with defaults. Both files are self-documented, allowing parameters to be inspected and edited before execution. Once these two parameter files are configured, *run_from_config*(sessions, params) executes the analysis identically to a GUI run and also executes a series of runs if its input is a list of parameters files. As introduced earlier, *run_from_config*(combined) additionally supports exact replay from a single self-contained config file generated by a previous run. Full installation instructions, quickstart guide and headless run step by step workflow are available in the online documentation (https://cicada-analysis.readthedocs.io/en/latest/analysis/headless_workflow/).

**Figure 2.**
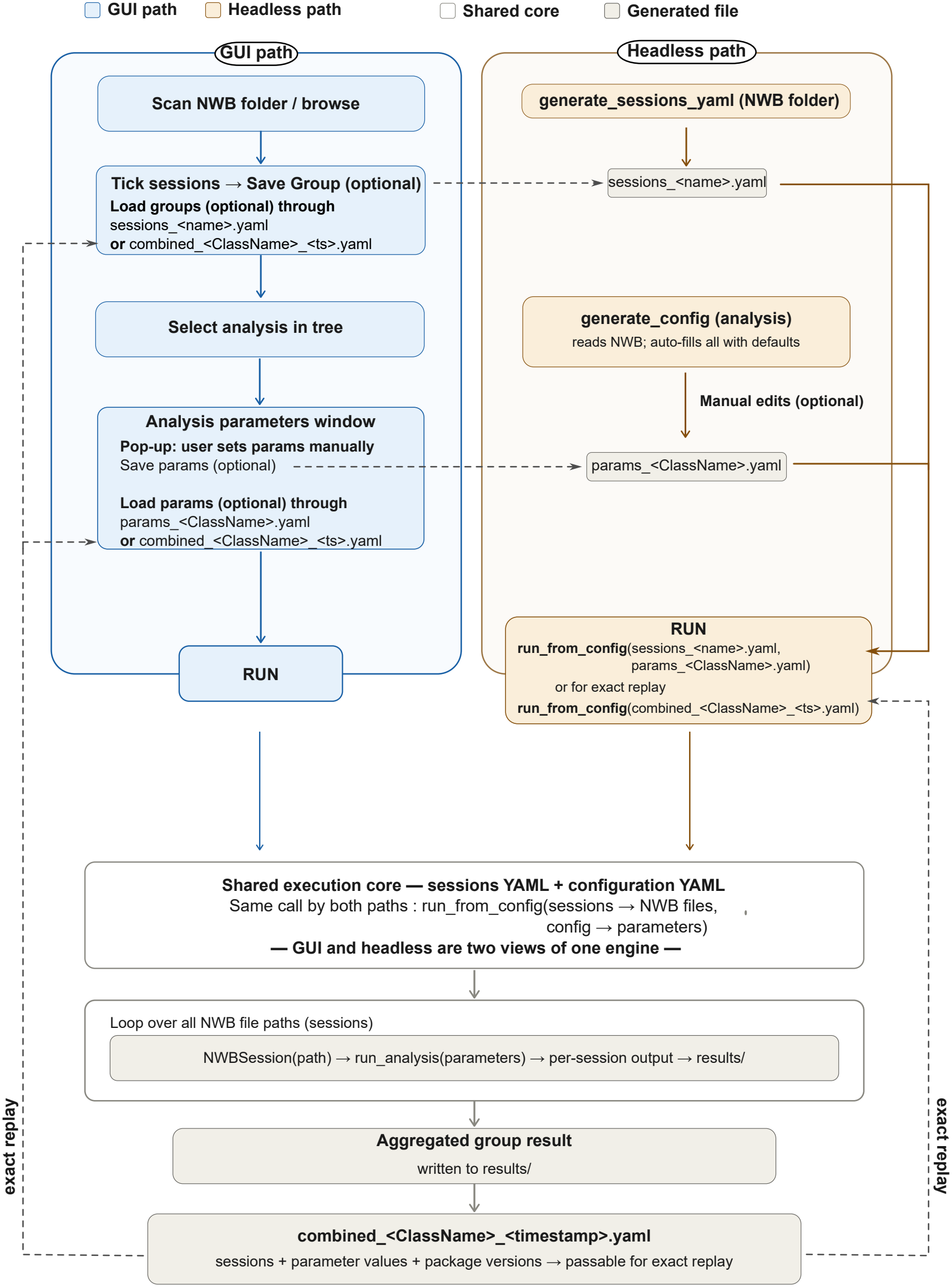
GUI and headless execution paths converge on a shared analysis core. Both the GUI path (blue) and the headless path (orange) ultimately call the same *run_analysis*(), ensuring strict equivalence between the two modes. In the GUI path, sessions are selected interactively and parameters set via the analysis parameters app. In the headless path, a session YAML is passed to *generate_config*(), which auto-populates a params_<ClassName>.yaml that the user then edits before calling *run_from_config*(sessions, params). In both cases, execution loops over all NWB files, calling *NWBSession*(path) and *run_analysis*() per session, and writes aggregated results to *results/*. Upon completion, a combined_<ClassName>_<timestamp>.yaml encoding sessions, parameter values, and package versions is generated, enabling exact replay via *run_from_config*(combined) or via the *load* options in the gui.

### R3. Demonstration use cases of end-to-end existing plugins

To illustrate the range of analyses accessible through the 16 core plugins of ***cicada-analysis***, we present five use cases (Figure 3). These examples are illustrative rather than exhaustive; a full description of all available plugins is provided in the online documentation (https://cicada-analysis.readthedocs.io/en/latest/analysis/cicada_analysis_plugins/). All analyses shown can be run from ***cicada-gui***, and are equally accessible through the headless runner as described earlier. Most of these analyses were done on demo NWBs provided with the pipeline, which consist of 2-minute excerpts of public DANDI datasets and of one unpublished dataset. We first demonstrate three analyses applied to two-photon calcium imaging data (Figure 3 - Panels A1-A2). In A1 we reused the already published in vivo 2-photon imaging data from the CA1 region of the hippocampus in mouse pups (DANDI 000219). We used the *cell map* plugin, which produces a contour map of detected cells color-coded by cell types, alongside a spike raster (*display activity* plugin) inferred from calcium traces and a connectivity graph in which nodes are color-coded by cell type and edges reflect functional connectivity (*connectivity graph* plugin). In A2 (DANDI 000728), the same *cell map* plugin is used with transient frequency coding, paired with the *display activity* plugin showing fluorescence traces with overlaid vertical markers at sensory stimulus times, and the *single cell PSTH* plugin showing single-trial and averaged responses of an individual cell to a given stimulus.

**Figure 3.**
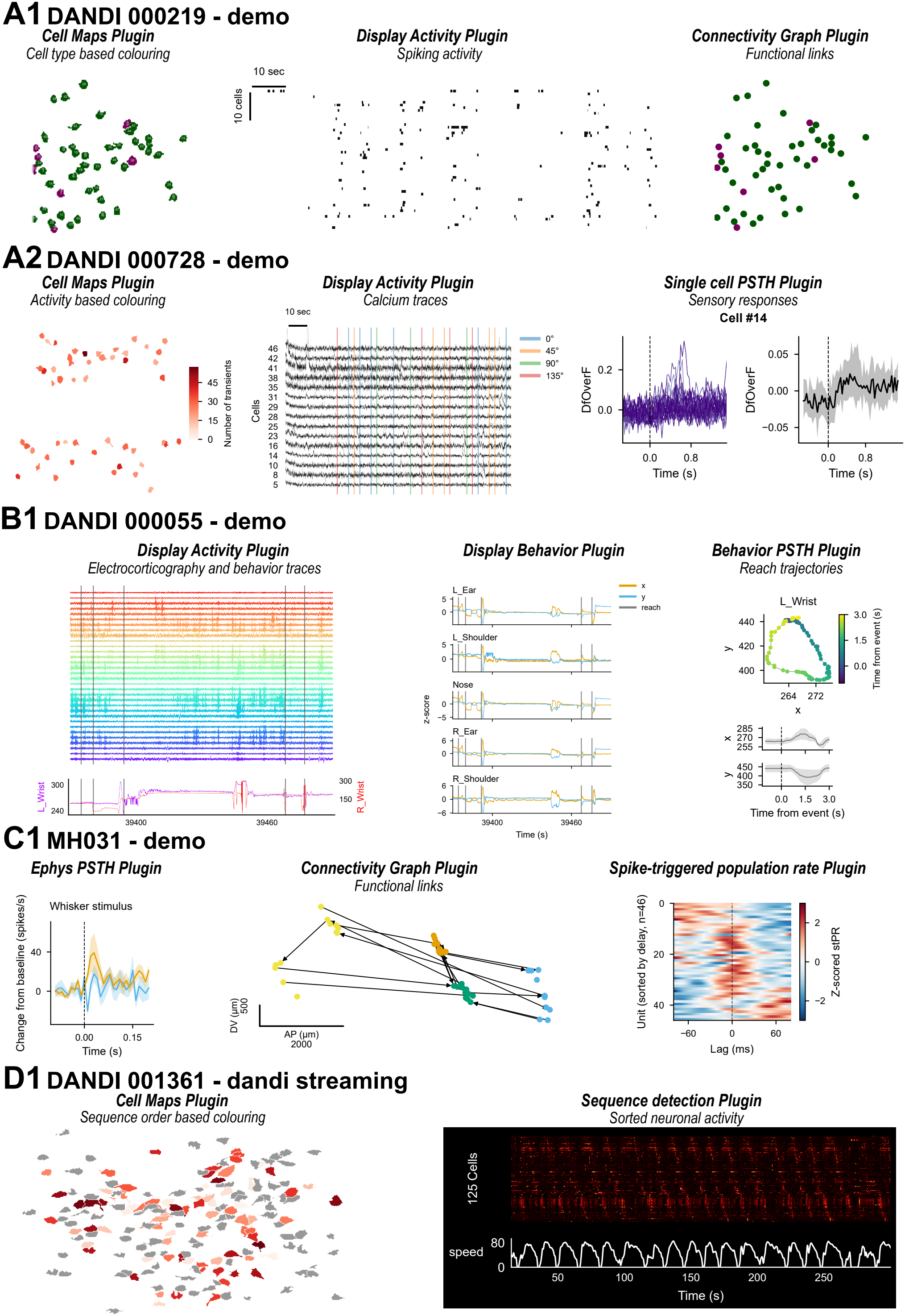
Example use cases across various neurophysiology data modalities. **(A1)** Two-photon calcium imaging dataset in the CA1 region of the hippocampus of a mouse pup. Left: cell contour map from the imaging field of view color-coded by cell types (pyramidal cells in green, interneurons in magenta). Middle: spike raster inferred from calcium traces. Right: connectivity graph with nodes color-coded by cell type. **(A2)** Two-photon calcium imaging dataset of primary visual cortex. Left: cell contour map of the imaging field of view, coloured by transient frequency. Middle: fluorescence traces with overlaid visual stimulus times. Right: single-cell PSTH showing single-trial and averaged responses to all visual stimuli. **(B1)** Human electrocorticography dataset. Left: Electrocorticography and behavior traces (wrist position) with overlaid reach times. Middle: Behavior traces with overlaid reach times. Right: Top: Average reach trajectory of the wrist in 2-d space. Bottom: PSTH of those trajectories. **(C1)** Large-scale extracellular electrophysiology dataset. Left: Baseline-subtracted PSTH showing neuronal responses to a single C2 whisker deflection averaged per area (primary whisker somatosensory cortex (orange) and midbrain (blue)). Middle: Connectivity graph with primary whisker somatosensory whisker (orange), midbrain (blue), thalamus (green) and motor areas (yellow) in a coronal projection of cells. Lines indicate putative monosynaptic connection. Right: Population coupling strength of individual neurons ordered by their coupling delay. **(D1)** Two-photon calcium imaging dataset in the CA1 region of the hippocampus of the adult mouse. Left: cell contour map of the imaging field of view color-coded by order of recruitment within a recurrent activity sequence detected in VR navigating mice; non-participating cells in grey. Right: corresponding sequence heatmap (top) along with speed of the mouse (bottom).

In Figure 3 - Panel B1 (DANDI 000055), the *display activity* plugin now shows simultaneous electrocorticography and behavior traces overlaid with vertical markers of reach initiations, and we then illustrate behavior only analyses, with the *display behavior* plugin showing different behavior traces within the same snapshot of time, paired with the *behavior psth* plugin showing the average reach trajectories. We then demonstrate analyses applied to single-unit electrophysiological data (Figure 3 - Panel C1) using a large-scale extracellular recording dataset (Unpublished - Bisi, Hamon). We used the *electrophysiology PSTH* plugin to show baseline-subtracted neuronal activity following a single whisker stimulus averaged per area. We then used the *Connectivity graph* plugin which infers putative monosynaptic connections using a previously described method (English et al., 2017) and plots it in a 2D projection of atlas space. On the same recording the plugin *Spike-triggered population rate* was run, computing for each unit a coupling strength and delay to the population as recently defined (Bimbard et al., 2026). Finally, we show analyses done through direct DANDI streaming of another two-photon imaging dataset (DANDI 001361) and during which we used the *sequences detection* plugin based on previously described method (Villette et al., 2015) to identify recurring activity patterns of hippocampal CA1 neurons; the contour map colors cells by their order of recruitment within the detected sequence, with non-participating cells shown in grey (Figure 3 - Panel D1). Notably, A1, A2 and D1 use the same *cell map* plugin and A1, A2 and B1 use the same *display activity* plugin, demonstrating how different parameter files alone produce outputs tailored to distinct scientific questions, without modifying any code.

Together these examples illustrate that CICADA supports analyses ranging from simple visualization to multi-step pattern detection. They also highlight that the same plugin, run with different parameter files, can address distinct scientific questions without modifying any code, whether executed through ***cicada-gui*** or ***cicada-analysis*** headless runner.

### R4. cicada-analysis — Community extension through plugins and lab packages

The CICADA pipeline is built to grow through community contributions. We designed it to make it easy for a laboratory to turn analysis code that would otherwise stay as a lab-specific NWB script into a robust, documented and shareable *cicada-analysis* plugin. A *cicada-analysis* plugin is a Python class that inherits from the *CicadaAnalysis* parent class and follows a fixed skeleton (Figure 4). The three class methods to implement each have a well-defined role. First, *check_data()* declares the data the plugin needs. It is called when sessions are loaded to filter the analysis tree, so only compatible plugins are shown for a given dataset. Second, *setup_parameters()* declares the parameter schema, which drives both the GUI parametrization window and YAML config generation. Third, *run_analysis()* contains the analysis logic. It receives the same parameter set whether the run was started from the GUI or in headless mode. The plugin also declares a *family_id*, which sets the family name under which it appears in the analysis tree. This fixed structure lets the framework handle discovery, parametrization, execution and provenance in the same way for every plugin, whether the core team or an external laboratory wrote it.

**Figure 4.**
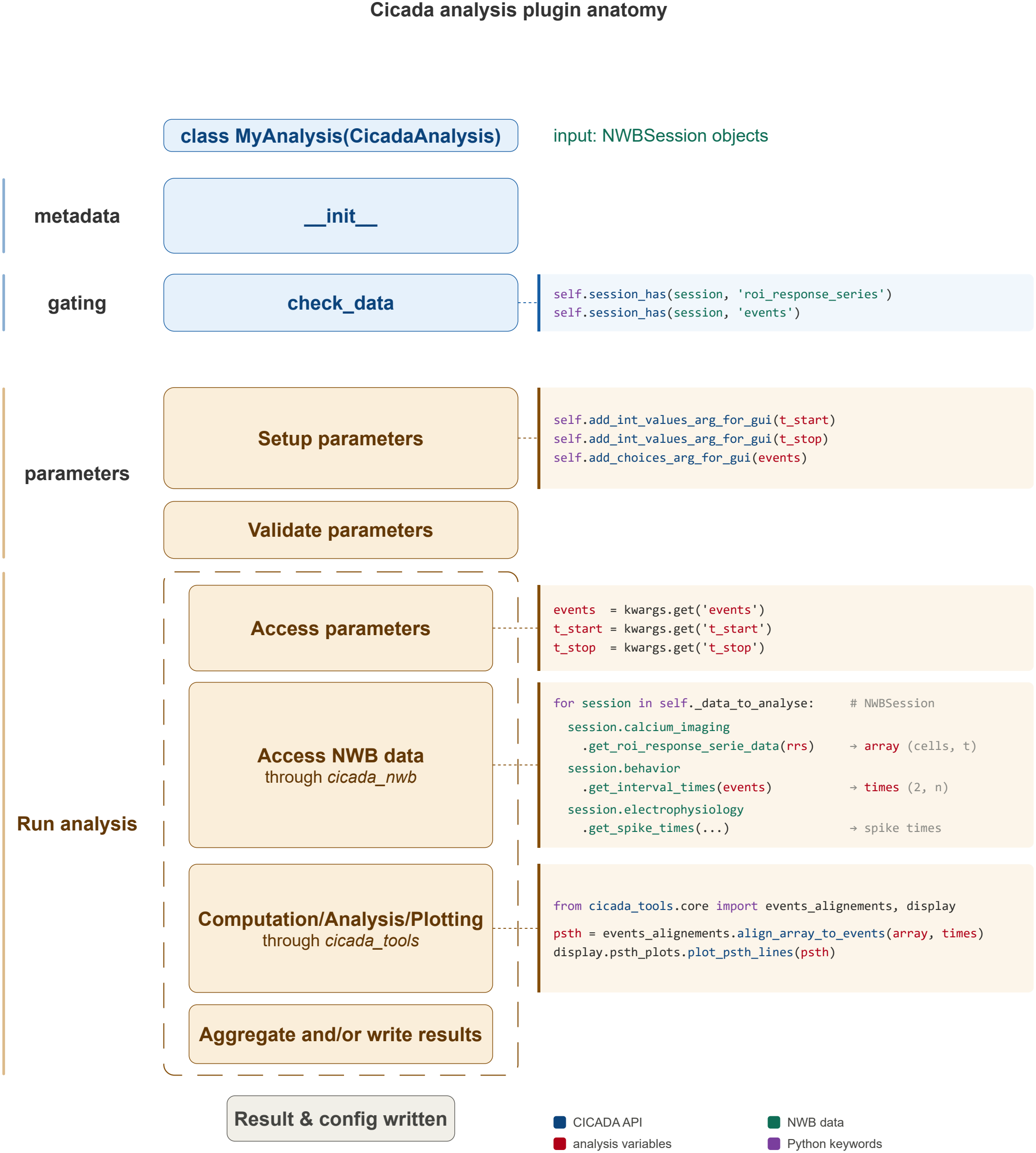
‘cicada-analysis’: plugin anatomy. A plugin is a Python class inheriting from *CicadaAnalysis* parent class and organized into four sequential components. The_init_method (metadata, blue) declares the plugin name, description, and *family_id. check_data*() (gating, blue) is called when sessions are loaded to determine whether the plugin is compatible with the loaded sessions, controlling its visibility in the analysis tree. *setup_parameters*() (parameters, orange) declares the parameter schema that drives both the GUI parametrization window and YAML config generation. *run_analysis*() (execution, orange) retrieves NWB data via ***cicada-nwb*** (standard submodules or the lab submodule *session*.*<lab>*), calls *cicada-tools* or lab tool functions for computation, and aggregates and writes results. Upon completion, results and a combined configuration file are written to disk.

Writing a plugin starts from a provided template that runs as is (https://cicada-analysis.readthedocs.io/en/latest/overview/developers/lab/#writing-an-analysis-plugin). The developer renames the class and implements the three methods, delegating computation to *cicada-tools* functions. During development, the plugin file can sit in a plain folder passed to CICADA (from the GUI or as a runner argument), where it is picked up at once. Once it works, it moves into a lab package: an ordinary Python package installed next to the core packages. A lab package has three parts and can be built from the lab package template available in the pipeline for any laboratory that wants to build its own (https://cicada-analysis.readthedocs.io/en/latest/overview/developers/lab/#writing-a-lab-package). First, a lab submodule holds the lab’s NWB conventions: it extends ***cicada-nwb*** with methods that read the lab’s own NWB fields or extensions, and is attached to every session as *session*.*<lab>*, next to the core submodules. Second, lab analysis tools, like cicada-tools, receive arrays and tables and never read NWB files. Third, lab plugins extend the core plugin collection, using the lab submodule, the lab tools and the shared cicada-tools. The lab package registers its parts through two standard Python entry points, declared in its pyproject.toml (cicada_nwb.lab_submodules and cicada_analysis.plugins). Once the package is installed with pip, the lab submodule extending ***cicada-nwb*** is available on every *NWBSession(path)* and the lab plugins are made accessible in the GUI analysis tree and through the headless runner. No file of the core framework is modified, forked or copied.

Distributing lab-specific CICADA extensions as packages makes them robust as well as shareable. First, it facilitates version checks since each lab package declares the range of core versions it was written for and CICADA checks this range when loading the package. A lab package written for another core version is not loaded, and a message gives the reason, instead of the analysis failing later. Each run also writes the full list of installed packages next to its combined YAML, including every lab package with its version and, for a git install, its commit, so a result can be traced to the exact lab code that produced it. Second, it allows laboratories to keep their code in their own repository, public or private, releasing it independently of the core pipeline. The core packages stay limited to analyses that use standard NWB types only. We applied this model to our own laboratories. The plugins previously bundled in ***cicada-analysis*** that rely on lab-specific NWB content or that address lab-specific analysis needs now live in two lab packages, cicada-cossart (14 plugins) and cicada-petersen (11 plugins). They are discovered through the same entry points as any third-party package, while the 16 core plugins remain in ***cicada-analysis***. Lab packages are also the route into the core collection. A lab plugin that relies only on standard NWB types can be proposed for transfer to ***cicada-analysis***, together with the tools it uses. Before joining the core, it is run on every demo NWB file with default and randomly drawn parameter sets, and the numbers it saves are compared with stored references on each change, so that a plugin’s output cannot change unnoticed. Writing a lab plugin to the same conventions from the start (standard NWB types read through ***cicada-nwb***, computation in tools, declared parameters, seeded randomness) makes this transfer a matter of moving files.

In this model, ***cicada-nwb*** and *cicada-tools* are the contributors’ entry point into CICADA. ***cicada-nwb*** is the layer through which every plugin reads data. A single entry point, *NWBSession(path)*, opens a local NWB file or a DANDI asset (streamed, or downloaded to a local cache) in read-only mode, and exposes its content through thematic submodules: *session*.*metadata, session*.*calcium_imaging, session*.*electrophysiology, session*.*behavior, session*.*widefield* and *session*.*optogenetic* (https://cicada-analysis.readthedocs.io/en/latest/nwb/). Its getters return standard Python objects (numpy arrays, pandas dataframes, lists, dictionaries) and load data lazily, reading only the requested slices, which keeps gigabyte-scale and streamed files practical. Plugins never address NWB paths directly: they ask the session what a file contains (for example which ROI response series are available, or whether a series holds continuous traces or a binary raster) and request the data through these methods. The same plugin therefore runs on files from different laboratories that store the same standard NWB type in different places. Lab submodules follow the same model for non-standard content and appear on every session as *session*.*<lab>*.

*cicada-tools* gathers the computation shared by all plugins: more than 250 documented functions in 12 thematic submodules (core, imaging, ephys, widefield, behavior, graphs, display, stats, sequences, connectivity, clustering and cell assemblies). They cover the operations most analyses need: event alignment, PSTH construction, behavioral analysis, processing of imaging and electrophysiological data, statistical testing (https://cicada-analysis.readthedocs.io/en/latest/analysis/cicada_tools/) and can also easily be extended through the lab package. Because these functions take standard Python objects (arrays, dataframes, lists) and never read NWB files, each one can be unit-tested on its own and reused across plugins. Both packages can also be imported directly into a notebook, for example to explore a dataset or prototype a computation before writing a plugin. Their main role in CICADA, however, is to supply the building blocks of plugins, so that new analyses are written once, gain GUI exposure, YAML parametrization, multi-session support and provenance tracking at no extra cost.

## CONCLUDING REMARKS

The purpose of CICADA is not only to offer a versatile set of pre-defined analysis pipelines, but also to provide a general framework for organizing, standardizing, and documenting analysis workflows. With its multiple entry points, CICADA intends to be easily accessible to all users, flexible, and scalable. Also of importance, CICADA is designed to cover a wide range of experimental modalities, from behavior to electrophysiology and calcium imaging.

A researcher unfamiliar with Python can run existing analyses entirely through ***cicada-gui*** without writing any code, or through the ***cicada-analysis*** headless runner with minimal scripting. A developer can write a plugin on top of ***cicada-nwb*** and *cicada-tools* that immediately gains GUI exposure, YAML parametrization and multi-session support, and share it, together with the lab’s NWB conventions and tools, as a lab package that any CICADA installation discovers once installed. A laboratory’s non-standard NWB content is read through a lab submodule in the same package, without changing the core. Each level is independently useful and adopting a higher level of integration is always optional.

One limitation is that extending ***cicada-nwb*** to support non-standard NWB fields via the lab submodule system requires familiarity with PyNWB, which may represent a barrier for some laboratories. However, the lab package template, with one working example of each part and its tests, is meant to lower this barrier. Two lab packages exist today (cicada-cossart, cicada-petersen); the model is open to any laboratory, and lab packages are the path by which new analyses reach the core collection.

Beyond its architectural flexibility, CICADA addresses three practical needs that are often unmet by lab-specific analysis scripts. First, it facilitates access to robust, parameterized analysis workflows that non-expert users can run without modifying code. Second, it ensures the reproducibility of the analyses and supports the long-term preservation of analysis methods within a laboratory, through YAML-serialized configurations that fully document how any result was produced. Third, it facilitates the sharing of analysis workflows across laboratories. A lab plugin written to the CICADA conventions is shared by installing its lab package. Once it relies only on standard NWB types, it can be proposed for the core collection, where it runs on any NWB file that contains the data it needs. Future development will be driven by community contributions: new lab packages, and the transfer of their plugins into the core collection.

## Acknowledgements

ACKNOWLEDGEMENTS

We thank all members of Rosa Cossart and Carl Petersen laboratories for their comments on the pipeline and on the manuscript.

## CODE AVAILABILITY

- CICADA pipeline source code:
  - ***cicada-gui***: https://gitlab.com/cossartlab/cicada_gui
  - ***cicada-analysis***: https://gitlab.com/cossartlab/cicada_analysis
  - ***cicada-nwb***: https://gitlab.com/cossartlab/cicada_nwb Documentation: https://cicada-analysis.readthedocs.io Released code: https://pypi.org/project/cicada-analysis/
- CICADA lab packages
  - ***cicada-cossart***: https://gitlab.com/cossartlab/cicada_cossart
  - ***cicada-petersen***: https://github.com/LSENS-BMI-EPFL/cicada_petersen

## DATASETS - DANDI Archive

**000713:** Allen Institute (2024). Allen Institute - Visual Behavior - Neuropixels (Version 0.240702.1725). DANDI Archive. https://doi.org/10.48324/dandi.000713/0.240702.1725

**000728:** Allen Institute (2024). Allen Institute - Visual Coding - Optical Physiology (Version 0.240827.1809). DANDI Archive. https://doi.org/10.48324/dandi.000728/0.240827.1809

**000219:** Dard, R., Picardo, M., Cossart, R. (2026). Two photon calcium imaging in the CA1 region of the hippocampus in neonatal mice (Version 0.260826.1155). DANDI Archive. https://doi.org/10.48324/dandi.000219/0.260826.1155

**000044:** Grosmark, A. D., Buzsáki, G. (2026). Diversity in neural firing dynamics supports both rigid and learned hippocampal sequences (Version 0.261005.2101). DANDI Archive. https://doi.org/10.48324/dandi.000044/0.261005.2101

**000055:** Peterson, S. M., Singh, S. H., Dichter, B., Scheid, M., Rao, R. P. N., Brunton, B. W. (2022). AJILE12: Long-term naturalistic human intracranial neural recordings and pose (Version 0.220127.0436). DANDI Archive. https://doi.org/10.48324/dandi.000055/0.220127.0436

**001361:** Sosa, M., Plitt, M. H., Giocomo, L. M. (2025). A flexible hippocampal population code for experience relative to reward (Version 0.251124.0550). DANDI Archive. https://doi.org/10.48324/dandi.001361/0.251124.0550 **000409:** International Brain Laboratory et al. IBL - Brain Wide Map (Version 0.260309.1324). DANDI Archive. https://doi.org/10.48324/dandi.000409/0.260309.1324

**001847:** Bech, P., Dard, R., Lebert, J., Smith, L., Bisi, A., Renard, A., Crochet, S., Petersen, C. (2026). Retrosplenial cortex enables context-dependent goal- directed sensorimotor transformation (Version 0.260610.1400). DANDI Archive. https://doi.org/10.48324/dandi.001847/0.260610.1400

## DATASETS - Unpublished

**MH031:** unpublished data of the Petersen lab (Laboratory of Sensory Processing, EPFL), shared with the lab’s permission for the CICADA demo and tests.

## REFERENCES

Bech P, Dard RF, Lebert J, Smith L, Bisi A, Renard A, Crochet S, Petersen CC. 2026. Retrosplenial cortex enables context-dependent goal-directed sensorimotor transformation. eLife 14. DOI: 10.7554/eLife.109717.1

Bimbard C, Harris KD, Carandini M. 2026. Invariant activity sequences across the mouse brain. DOI: 10.64898/2025.12.20.695676

Bocchio M, Gouny C, Angulo-Garcia D, Toulat T, Tressard T, Quiroli E, Baude A, Cossart R. 2020. Hippocampal hub neurons maintain distinct connectivity throughout their lifetime. Nature Communications 11:4559. DOI: 10.1038/s41467-020-18432-6

Buccino AP, Hurwitz CL, Garcia S, Magland J, Siegle JH, Hurwitz R, Hennig MH. 2020. SpikeInterface, a unified framework for spike sorting. eLife 9:e61834. DOI: 10.7554/eLife.61834, PMID: 33170122

Chung JE, Magland JF, Barnett AH, Tolosa VM, Tooker AC, Lee KY, Shah KG, Felix SH, Frank LM, Greengard LF. 2017. A Fully Automated Approach to Spike Sorting. Neuron 95:1381–1394.e6. DOI: 10.1016/j.neuron.2017.08.030

Cossart Lab / CICADA. 2019. GitLab. https://gitlab.com/cossartlab/cicadaDANDI. n.d. DANDI. https://about.dandiarchive.org/

Dard RF, Leprince E, Denis J, Rao Balappa S, Suchkov D, Boyce R, Lopez C, Giorgi-Kurz M, Szwagier T, Dumont T, Rouault H, Minlebaev M, Baude A, Cossart R, Picardo MA. 2022. The rapid developmental rise of somatic inhibition disengages hippocampal dynamics from self-motion. eLife 11:e78116. DOI: 10.7554/eLife.78116

English DF, McKenzie S, Evans T, Kim K, Yoon E, Buzsáki G. 2017. Pyramidal Cell-Interneuron Circuit Architecture and Dynamics in Hippocampal Networks. Neuron 96:505–520.e7. DOI: 10.1016/j.neuron.2017.09.033, PMID: 29024669

Flynn GM, M A, Baker C, Ngo T, Clark J, fairdataihub-bot Portillo D, Patel B, Ly R, Soundarajan S, Srivastava H, Bhansali I, Dichter B, Helmer K, Tauffer L, Adkisson-Floro P, Gillespie T, Buccino A, Magland J, Gunalan K dhmjhu. 2026. NeurodataWithoutBorders/nwb-guide: 1.1.0. DOI: 10.5281/zenodo.18930548

Giovannucci A, Friedrich J, Gunn P, Kalfon J, Brown BL, Koay SA, Taxidis J, Najafi F, Gauthier JL, Zhou P, Khakh BS, Tank DW, Chklovskii DB, Pnevmatikakis EA. 2019. CaImAn an open source tool for scalable calcium imaging data analysis. eLife 8:e38173. DOI: 10.7554/eLife.38173

Lee KH, Denovellis EL, Ly R, Magland J, Soules J, Comrie AE, Gramling DP, Guidera JA, Nevers R, Adenekan P, Brozdowski C, Bray SR, Monroe E, Bak JH, Coulter ME, Sun X, Broyles E, Shin D, Chiang S, Holobetz C, Tritt A, Rübel O, Nguyen T, Yatsenko D, Chu J, Kemere C, Garcia S, Buccino A, Frank LM. 2024. Spyglass: a framework for reproducible and shareable neuroscience research. DOI: 10.1101/2024.01.25.577295

Leprince E, Dard RF, Mortet S, Filippi C, Giorgi-Kurz M, Bourboulou R, Lenck-Santini P-P, Picardo MA, Bocchio M, Baude A, Cossart R. 2023. Extrinsic control of the early postnatal CA1 hippocampal circuits. Neuron S0896627322010868. DOI: 10.1016/j.neuron.2022.12.013

Leprince E, Filippi C, Mantez M, Dard RF, Dichio V, Majnik J, Cretella M, Bocchio M, Picardo MA, Monasson R, Platel J-C, Cossart R. 2026. Distinct developmental trajectories of externally and internally generated hippocampal sequences and assemblies. Current Biology. DOI: 10.1016/j.cub.2026.04.053

Mayorquin H, Baker C, Adkisson-Floro P, Weigl S, Trappani A, Tauffer L, Rübel O, Dichter B. 2025. NeuroConv: Streamlining Neurophysiology Data Conversion to the NWB Standard. Python in Science Conference, 2025 245–258. DOI: 10.25080/cehj4257

NWBWidgets. n.d. https://github.com/NeurodataWithoutBorders/nwbwidgets?tab=readme-ov-file

Open Source Brain. n.d. https://v2.opensourcebrain.org/

Pachitariu M, Sridhar S, Pennington J, Stringer C. 2024. Spike sorting with Kilosort4. Nature Methods 21:914–921. DOI: 10.1038/s41592-024-02232-7

Pachitariu M, Stringer C, Dipoppa M, Schröder S, Rossi LF, Dalgleish H, Carandini M, Harris KD. 2017. Suite2p: beyond 10,000 neurons with standard two-photon microscopy. DOI: 10.1101/061507

Ratsifandrihamanana MR, Dard RF, Denis J, Cossart R, Picardo MA. 2023. Protocol to image and analyze hippocampal network dynamics in non-anesthetized mouse pups. STAR Protocols 4:102760. DOI: 10.1016/j.xpro.2023.102760

Rübel O, Tritt A, Ly R, Dichter BK, Ghosh S, Niu L, Baker P, Soltesz I, Ng L, Svoboda K, Frank L, Bouchard KE. 2022. The Neurodata Without Borders ecosystem for neurophysiological data science. eLife 11:e78362. DOI: 10.7554/eLife.78362

Viejo G, Levenstein D, Carrasco SS, Mehrotra D, Mahallati S, Vite GR, Denny H, Sjulson L, Battaglia FP, Peyrache A. 2023. Pynapple, a toolbox for data analysis in neuroscience. eLife 12. DOI: 10.7554/eLife.85786.2

Villette V, Malvache A, Tressard T, Dupuy N, Cossart R. 2015. Internally Recurring Hippocampal Sequences as a Population Template of Spatiotemporal Information. Neuron 88:357–366. DOI: 10.1016/j.neuron.2015.09.052, PMID: 26494280

Yamane Y, Li Y, Matsumoto K, Kanai R, Desforges M, Gutierrez CE, Doya K. 2025. Optical Neuroimage Studio (OptiNiSt): Intuitive, scalable, extendable framework for optical neuroimage data analysis. PLOS Computational Biology 21:e1013087. DOI: 10.1371/journal.pcbi.1013087

Yger P, Spampinato GL, Esposito E, Lefebvre B, Deny S, Gardella C, Stimberg M, Jetter F, Zeck G, Picaud S, Duebel J, Marre O. 2018. A spike sorting toolbox for up to thousands of electrodes validated with ground truth recordings in vitro and in vivo. eLife 7:e34518. DOI: 10.7554/eLife.34518

